# Expression variations in Ectodysplasin-A gene (*eda*) may contribute to morphological divergence of scales in Haplochromine cichlids

**DOI:** 10.1101/2021.08.25.457685

**Authors:** Maximilian Wagner, Sandra Bračun, Anna Duenser, Christian Sturmbauer, Wolfgang Gessl, Ehsan Pashay Ahi

## Abstract

**Background:** Elasmoid scales are one of the most common dermal appendages and can be found in almost all species of bony fish differing greatly in their shape. Whilst the genetic underpinnings behind elasmoid scale development have been investigated, not much is known about the mechanisms involved in the shaping of scales. To investigate the links between gene expression differences and morphological divergence, we inferred shape variation of scales from two different areas of the body (anterior and posterior) stemming from ten haplochromine cichlid species from different origins (Lake Tanganyika, Lake Malawi, Lake Victoria and riverine). Additionally, we investigated transcriptional differences of a set of genes known to be involved in scale development and morphogenesis in fish.

**Results:** We found that scales from the anterior and posterior part of the body strongly differ in their overall shape, and a separate look on scales from each body part revealed similar trajectories of shape differences considering the lake origin of single investigated species. Above all, nine as well as 11 out of 16 target genes showed expression differences between the lakes for the anterior and posterior dataset, respectively. Whereas in posterior scales four genes (*dlx5, eda, rankl* and *shh*) revealed significant correlations between expression and morphological differentiation, in anterior scales only one gene (*eda*) showed such a correlation. Furthermore, eda displayed the most significant expression difference between species of Lake Tanganyika and species of the other two younger lakes. Finally, we found genetic differences in downstream regions of *eda* gene (e.g. in the *eda*-*tnfsf13b* inter-genic region) that are associated with observed expression differences. This is reminiscent of a genetic difference in the *eda*-*tnfsf13b* inter-genic region which leads to gain or loss of armour plates in stickleback.

**Conclusion:** These findings provide evidence for cross-species transcriptional differences of an important morphogenetic factor, *eda*, which is involved in formation of ectodermal appendages. These expression differences appeared to be associated with morphological differences observed in the scales of haplochromine cichlids indicating potential role of eda mediated signal in divergent scale morphogenesis in fish.

## Background

Cichlids pose a great a model system for evolutionary biology, as they include some of the most striking examples of explosive speciation and adaptive radiation. Many aspects of their life history as well as their behaviour, coloration and feeding morphologies are well studied [1–3]. One of the most striking features is their repeated evolution of parallel eco-morphologies, especially across the radiations of the three East African Great Lakes, Lake Tanganyika (LT), Lake Malawi (LM) and Lake Victoria (LV) [4, 5]. These ecological adaptations are also the focus of many studies, as they promise the opportunity to shed light on different molecular mechanisms underlying repeated evolution and diversification [6, 7]. Regarding skeletal morphogenesis in particular the evolution of their jaws and their phenotypic plasticity are topics of ongoing research [7–12]. However, while the adaptive value of some of the investigated structures (e.g., feeding apparatus) can be more easily connected to certain ecological specializations [5, 13], this is not so obvious in others, such as scales.

Fish scales come in a vast array of different shapes and forms. As a part of the dermal skeleton, which amongst other structures also includes teeth, odontodes, spines and fin rays, these postcranial derivates evolved into morphologically and histologically diverse structures in Actinopterygii [14, 15]. Elasmoid scales, found in most of teleost species, form in the dermal mesenchyme and are mainly used for protection and hypothetically for hydrodynamic modifications [14, 16, 17]. While the elasmoid scales form relatively late in ontogeny and can take diverse forms, they share a composition consisting of three tissues, with elasmodin as the basal component formed in a characteristic plywood-like structure [15, 16]. Scale development, mostly studied in zebrafish, has been found to be orchestrated by several well-known pathways, including Hh, Fgf and Eda [16, 18–20], which are known to be also involved in the appendage formation across several vertebrate groups [21]. Mutations and allele variations in the Eda/Edar pathway, for example, have been linked to fish fin, scale and armour plate development as well as human and mouse hair and teeth growth [19, 22, 23]. Nevertheless, besides a recent extensive comparison of the scale morphology across

Lake Tanganyika cichlids [24], as well as a genetic study of scale shapes in two closely related Lake Malawi cichlids, which tied FgF signalling to scale shape variation [20], not much is known about the molecular mechanisms shaping the elasmoid scale.

In this study, we investigate the morphological differences in the anterior and posterior scales of 10 haplochromine cichlid fish species from three Great East African Lakes, i.e., Lake Tanganyika (LT), Lake Malawi (LM) and Lake Victoria (LV) as well as a riverine haplochromine cichlid species. After identification of a stably expressed reference gene, we also investigate transcriptional differences of a set of genes known to be involved in scale development and morphogenesis in fish. Finally, we tried to find links between the gene expression differences and morphological divergence in both anterior and posterior scales. Our results provide cross-species expression comparisons of scale related genes in haplochromine cichlids and implicate expression differences by which formation of distinct scale morphologies might be determined.

## Methods

### Fish husbandry and sampling

Ten haplochromine cichlid species; three species from Lake Tanganyika, four species from Lake Malawi, two species from Lake Victoria, and one riverine haplochromine species, were selected for this study (Fig. 1A). The fish were kept and raised in standardized tanks and rearing conditions with the same diet (Spirulina flakes) until they displayed mating behaviour. Between 5 to 11 adult females per species were sampled for morphological analysis and 4 adult females were sampled for gene expression investigation. The sampled fish species were sacrificed by euthanization in with 0.5 g MS-222/litre of water, and 5 anterior and posterior scales from left side of the body were removed for morphological analysis (Fig. 1B), whereas similar numbers of scales were taken from both sides and all anterior or posterior scales from each fish were pooled for gene expression part.

**Figure 1.**
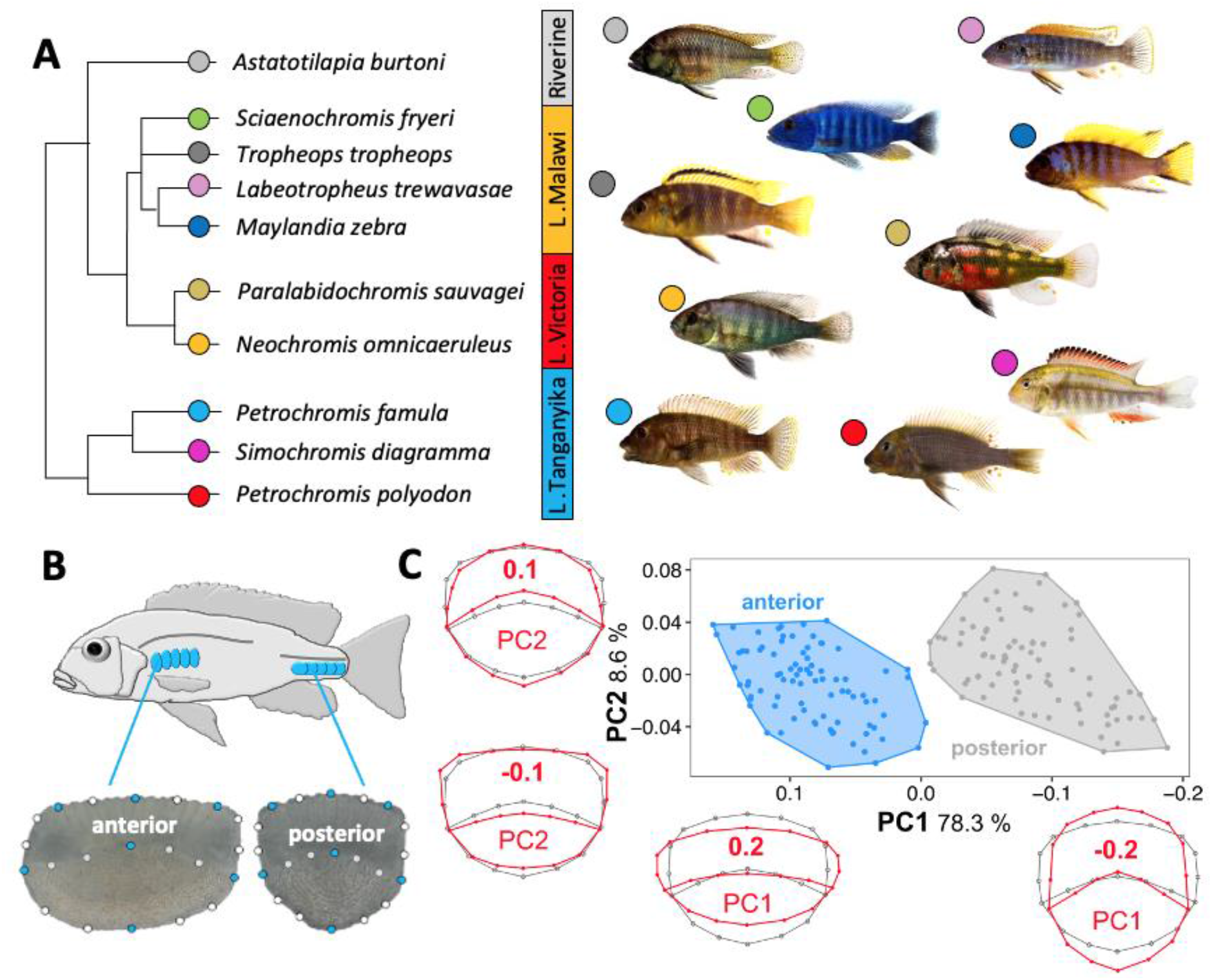
The haplochromine cichlid species and descriptions of the scale samples. (a) A simplified phylogenetic relatedness of the East African haplochromine cichlid species used in this study. (b) Positions of the anterior and posterior scales used in this study and landmarks used for the geometric morphometric analyses in both anterior (left) and posterior (right) scales shown as an example for *Petrochromis famula*. Blue dots represent major landmarks, white dots semi-landmarks and 1 mm scale bars are given below the images. (c) Principal component analysis (PCA) plots clearly separate scales from the anterior and posterior part of the body. Additional, warped outline drawings illustrate major shape changes along the axis (red) compared to the overall mean shape (grey).

### Morphological analysis

To infer shape differences of scales from divergent African cichlids from different lakes a 2D geometric morphometric framework was deployed. Due to major morphological differences of the scales they were separately investigated for the anterior and posterior part of the body (Fig. 1 B and C). Standardized images of scales were taken with a KEYENCE VHX-5000 digital microscope (KEYENCE Germany GmbH). 84 adult specimens from 10 cichlid species inhabiting the three major rift lakes which were reared under standardized aquarium conditions (*Astatotilapia burtoni* = 7; *Neochromis omnicaeruleus =* 10; *Petrochromis famula* = 11, *P. polyodon* = 7; *Paralabidochromis sauvage* = 5; *Simochromis diagramma* = 11; *Sciaenochromis fryeri* = 5; *Tropheops tropheops* = 9; *Labeotropheus trewavasae* = 9; Mz: *Maylandia zebra* = 10) were included for the geometric morphometric analyses. For each individual six scale replicates from the anterior and posterior part of the body were probed (Fig. 1B), leading to a total of 1.008 investigated scales. After randomizing pictures in tpsUtil v.1.6 (available at http://life.bio.sunysb.edu/morph/soft-utility.html), landmark digitization was conducted on a set of 7 fixed landmarks and 14 semi-landmarks (see Figure 1b for positions) in tpsDig v.2.26 (available at http://life.bio.sunysb.edu/morph/soft-utility.html). To ensure consistency, this step was conducted by a single investigator. Generalized Procrustes superimposition [25] was performed in tpsRelw v.1.65 (available at http://life.bio.sunysb.edu/morph/soft-utility.html) and aligned landmark configurations were exported for further analysis in MorphoJ v.1.06 [26]. In MorphoJ, single observations obtained from the six replicates were averaged to get the mean shape for each landmark. A Principal Component analysis (PCA) was applied to infer variation in morphospace among scale position (anterior vs. posterior), single specimen, and species. Subsequent analyses were based on separated datasets for anterior and posterior scale landmark setting, whereas PC-scores were exported for linear discriminant function analyses (LDA) in PAST v.4.1 [27]. To reduce the number of variables and control for putative over-separation of groups [28], only the first four principal components were used for the LDA. PCA and LDA plots were visualized in R v3.1.2 [29].

### RNA isolation and cDNA synthesis

As mentioned in the section above, 10 anterior and posterior scales from each fish were pooled for isolating the total RNA isolation in a single tube containing 0.25 mL of a tissue lysis buffer from Reliaprep RNA tissue miniprep system (Promega, #Z6111, USA) as well as one 1.4 mm ceramic bead to crush the scales. The scales were homogenized using a FastPrep-24 Instrument (MP Biomedicals, Santa Ana, CA, USA) and total RNA was extracted following the instructions provided by the manufacturer (adjusted protocol for small amounts of fibrous tissue). In summary, the instruction follows with mixing of the lysis buffer and homogenized scales with isopropanol and centrifuging the entire mix through a column provided by the kit, several RNA washing steps and a final DNase treatment step. The RNAs were quantified by a Nanophotometer (IMPLEN GmbH, Munich, Germany) and their quality was checked with RNA ScreenTapes on an Agilent 2200 TapeStation (Agilent Technologies). Next, the RNA samples with a RNA integrity number (RIN) above six were applied to first strand cDNA synthesis using 300ng of RNA and High Capacity cDNA Reverse Transcription kit (Applied Biosystems). The synthesized cDNA from each RNA sample was diluted 1:5 times in nuclease-free water to conduct qPCR.

### Gene selection, designing primers and binding site predictions

We selected eight candidate reference genes which have been frequently used in different studies of Haplochromine cichlids and have shown high expression levels in various connective tissues including skeletal tissues [10, 30–35]. Furthermore, we chose 16 target candidate genes, which are implicated in scale development and morphogenesis (Table 1). The primers were designed at conserved sequence of coding regions using the transcriptome data of several East African haplochromine species (*Astatotilapia burtoni, Aulonocara baenschi, Cyrtocara moorii, Pundamilia nyererei, Metriaclima zebra, Simochromis diagramma, Tropheus duboisi*, and *Gnathochromis pfefferi*) as well as two more distant species from different African cichlid tribes (*Oreochromis niloticus* and *Neolamprologus brichardi*) [7, 36–39]. The sequences from all the species were imported to CLC Genomic Workbench, version 7.5 (CLC Bio, Aarhus, Denmark), and after alignment, the exon/exon junctions were specified using the *Astatotilapia burtoni* annotated genome in the Ensembl database (http://www.ensembl.org) [40]. The primers were designed spanning exon junctions and a short amplicon size (<250 bp) as recommended to be optimal for qPCR quantification [41]. The primers were designed and assessed through Primer Express 3.0 (Applied Biosystems, CA, USA) and OligoAnalyzer 3.1 (Integrated DNA Technology) to minimize the occurrence of dimerization and secondary structures.

**Table 1.**
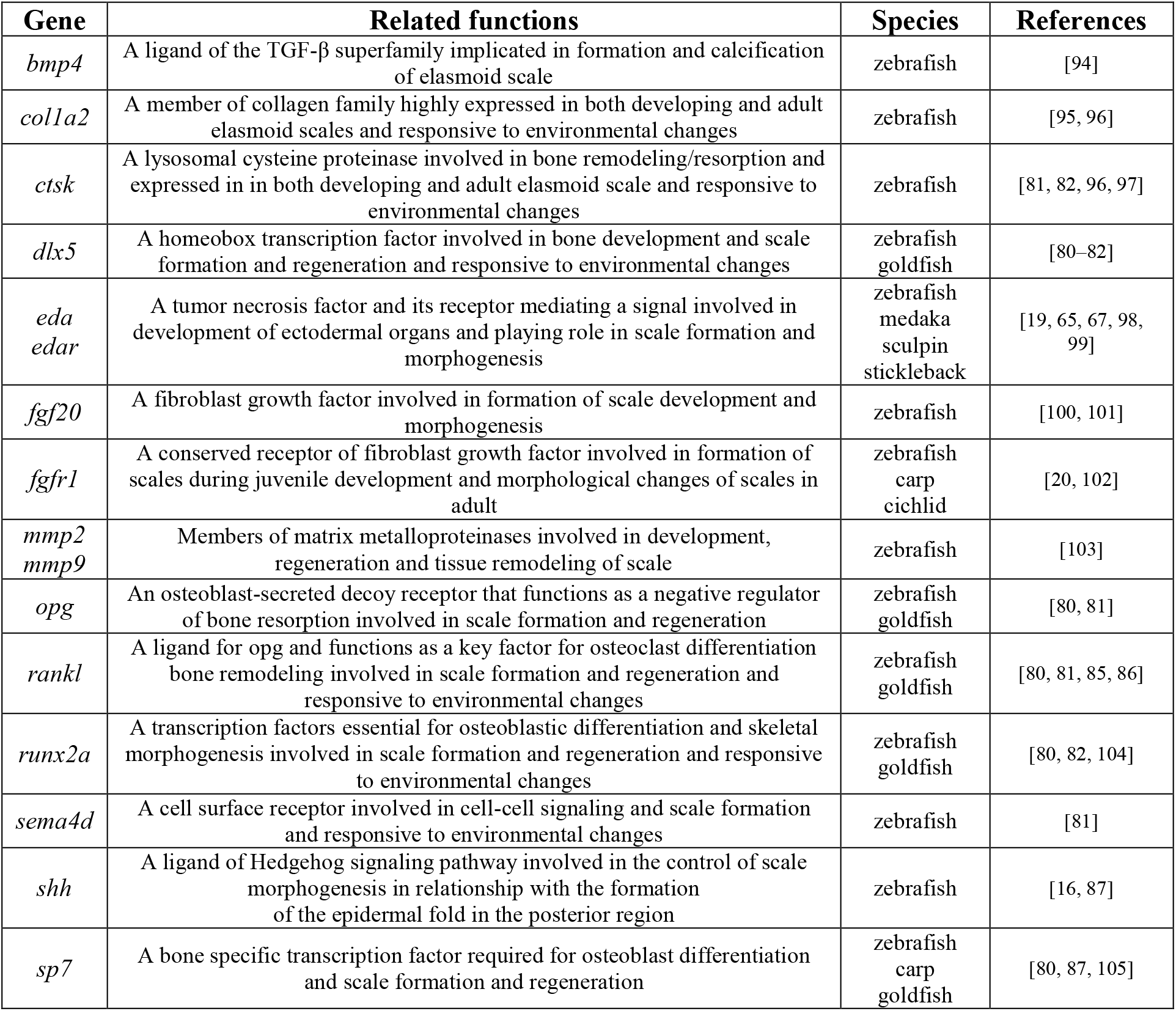
**Selected target genes involved in the development and/or morphogenesis of scales in teleost fish.**

We retrieved downstream sequences (3’UTR and inter-genic region) of *eda* gene for all the species in this study from European Nucleotide Archive (ENA) and Sequence Read Archive (SRA) in order to identify changes in potential binding sites. To do this, we used genomic sequences of the haplochromine species; *A. burtoni* (GCA_000239415.1), *P. famula* (GCA_015108095.1), *N. omnicaeruleus* (SRR12700904), *P. polyodon* (GCA_015103895.1), *S. fryeri* (ERX1818621), *S. diagramma* (GCA_900408965.1), *M. zebra* (GCA_000238955.1), *T. tropheops* (SAMEA2661272), *L. trewavasae* (SAMN12216683), and *P. sauvagei* (GCA_018403495.1). Next we identify the 3’UTR and inter-genic region of *eda* genes using the annotated genome of *A. burtoni* from Ensembl and aligned them using CLC Genomic Workbench. The different sequence motifs were identified and screened for potential TF binding sites using STAMP [42] and the PWMs obtained from the TRANSFAC database [43].

### qPCR and data analysis

The qPCR reactions were generated using Maxima SYBR Green/ROX qPCR Master Mix (2X) (Thermo Fisher Scientific, Germany) and the amplifications were conducted on ABI 7500 real-time PCR System (Applied Biosystems). The qPCR setups followed the recommended optimal sample maximization method [44]. The qPCR program, dissociation step and calculation of primer efficiencies were performed as described in our previous study [45] (Additional file 1).

Three different algorithms were applied to validate the most stable reference genes; BestKeeper [46], NormFinder [47] and geNorm [48]. The Cq value of the most stable reference gene was used as normalization factor (Cq _reference_) to calculate ΔCq of each target gene (ΔCq _target_ = Cq _target_ – Cq _reference_). The lowest expressed sample in each expression comparison was used as a calibrator sample and rest of the samples were subtracted from its ΔCq value to calculate ΔΔCq values (ΔCq _target_ – ΔCq _calibrator_). Relative expression quantities (RQ) were calculated through E^−ΔΔCq^ [49]. In order to perform statistical analysis, fold differences (FD) were calculated by transformation of RQ values to logarithmic values [50]. The significant expression differences were calculated using ANOVA statistical tests, followed by Tukey’s HSD *post hoc* tests. The correlations between gene expression and a morphometric parameter (canonical variate 1) were calculated through Pearson correlation coefficients (r) for each gene using R.

## Results

### Divergence in scale morphology

The principal component analysis (PCA) revealed a clear separation in overall average individual shape between anterior and posterior scales (Fig. 1D). PC1 and PC2 explained 78.3 % and 8.6 % of the total shape variation, respectively. Generally, on the first axis anterior scales are anterior-posteriorly more compressed compared to the posterior body part (see deformation grids in Fig. 1D). Along the second PC axis changes can be observed in the shape of the posterior scale field (narrow vs. wide), as well as in the lateral edges of the anterior scale field (edges vs. round).

While comparing different species, large variation in overall shape can be observed in the dataset which is restricted to anterior scales only. *Petrochromis famula, Maylandia zebra* and *Sciaenochromis fryeri* occupy large parts of the morphospace and overall, less intraspecific variation can be observed in other species (Fig. 2A). In the anterior dataset changes along the PC1 explain 47.1 % of total variation, and mainly affect the circularity of the overall shape (i.e., that scales get more compressed towards positive values). Changes along the second PC, which explains 19.6 % of the total shape variation, affect the posterior scale field (compression vs. expansion). Compared to anterior scales, less intraspecific shape variation can be observed in posterior scales, whereas PC1 explains 54.8 % and PC2 18.9 % of the total variation, respectively (Fig. 2B). PC1 separates two major clusters (*S. diagramma* + *P. famula* vs. rest) whereas changes along the axis mainly contribute to a dorso-ventrally versus anterior-posterior compression of the scale and the roundness of the anterior scale field. Along PC2 shape changes affect the expansion (or compression) of the anterior and posterior scale fields. Generally, the PCA only poorly resolves the lake (or phylogenetic) origin of the single species for both the anterior and posterior dataset.

**Figure 2.**
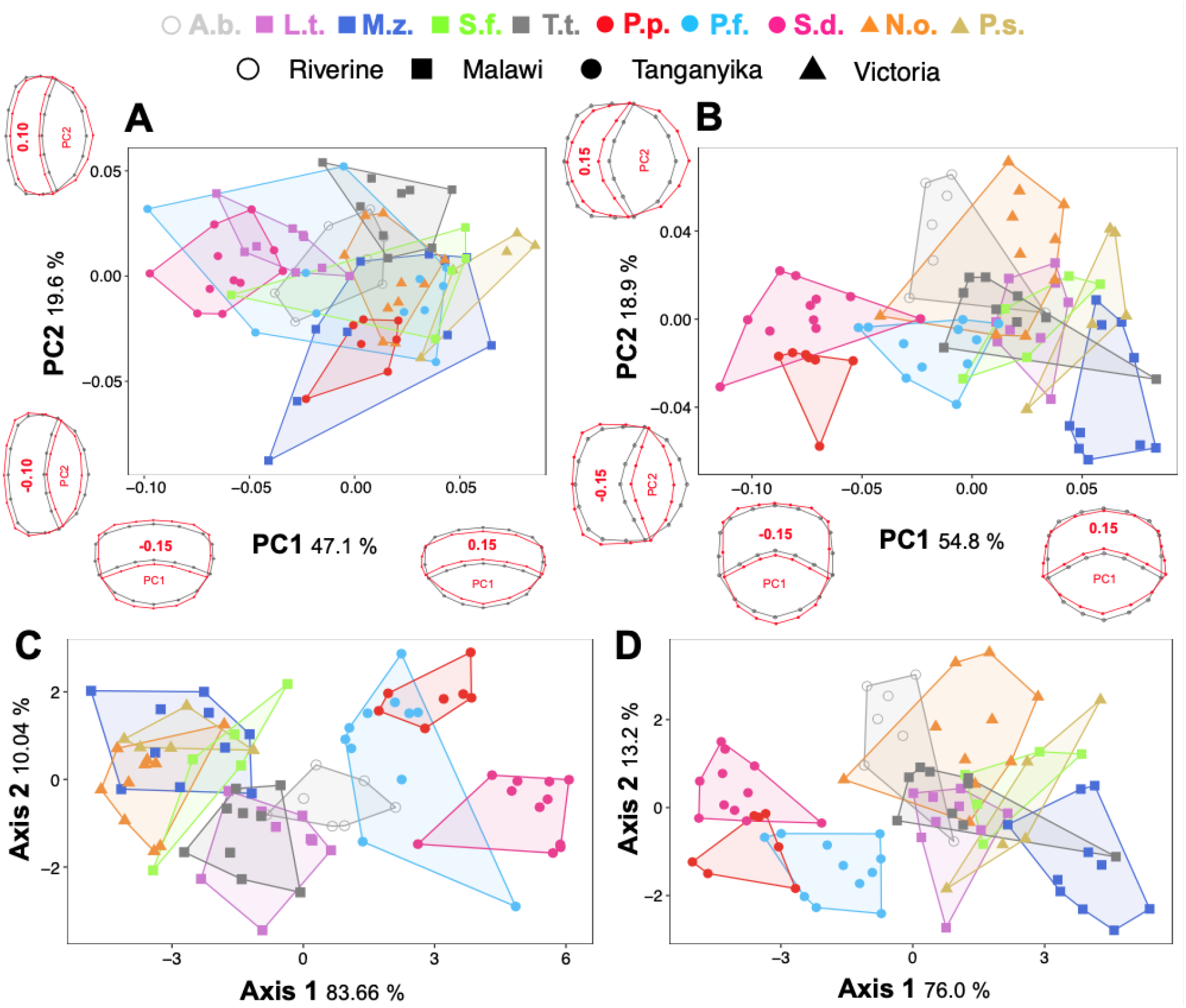
Morphospace of investigated scales from different species and body parts. Principal component analysis based on average shape of scales collected for the anterior (a) and posterior (b) part of the body and respective shape differences along the axis (grey: overall mean shape; red: shape change). Linear discriminant function analysis based on the first four PC-scores for anterior (c) and posterior (d) scales. All data points represent mean shapes obtained from 6 individually collected scales and shapes represent different lake origins. Abbreviations: A.b. Astatotilapia burtoni; N.o.: Neochromis omnicaeruleus; P.f.: Petrochromis famula, P.p.: P. polyodon; P.s.: Paralabidochromis sauvage S.d.: Simochromis diagramma; S.f.: Sciaenochromis fryeri; T.t.: Tropheops tropheops; L.t.: Labeotropheus trewavasae; M.z.: Maylandia zebra.

The linear discriminant function analysis (LDA) of anterior as well as posterior dataset correctly classified 77.38 % (jackknifed: 67.86 %) and 72.62 % (jackknifed: 65.48%) of species (Fig. 2C and D). The first axis explains 83.66 % and 76.00 % variance of the overall shape variability for the anterior and posterior dataset, respectively. In the anterior dataset, the first LD-axis separates three major clusters made up of samples from Lake Tanganyika, the riverine *Astatotilapia burtoni* and a joint Victoria-Malawi cluster. Similar results were obtained for the posterior dataset, whereas along the first axis the separation between the riverine *A. burtoni* and the Victoria-Malawi cluster is less prominent. Along the second axis, which explains 10.04 % and 13.2 % of the variance of the overall shape variability for the anterior and posterior dataset, respectively, mainly interspecific and intraspecific variation is portrayed. Overall, for the anterior dataset, 83.33% (jackknifed: 75 %) of species were correctly classified according to the lake origin, whereas single classification scores reached values of 100 % (jackknifed: 85.71 %) for *A. burtoni*, as well as 66.67 % (jackknifed: 57.58 %) for Malawi, 86.67 % (jackknifed: 86.67 %) for Victoria and 96.55 % (jackknifed: 86.21 %). In total, for the posterior dataset, 77.38 % (jackknifed: 70.24 %) of the individuals were correctly assigned to the lake origin, with 85.71 % (jackknifed: 71.43 %) for *A. burtoni*, as well as 66.67 % (jackknifed: 54.55 %) for Malawi, 53.33% (jackknifed: 46.67 %) for Victoria and 100 % (jackknifed: 100 %) for individuals from Tanganyika.

### Validation of stable reference genes

To quantify the expression levels of the selected target genes, the validation of stable reference gene(s) with least variation in expression across the anterior and posterior scales of different species is a necessary step [51]. The 8 candidates were selected from frequently used reference genes in studies of different tissues in East African cichlids [10, 30–35]. The candidate reference genes showed variable expression levels in the scales, and from highest to lowest expressed were respectively; *actb1, hsp90a, rps11, rps18, hprt1, gapdh, elf1a* and *tbp*. Interestingly, in both anterior and posterior scales, all the three software ranked *actb1* as the most stable reference gene with lowest expression variation across the cichlid species in this study (Table 2). Thus, we used the Cq value of *actb1* as normalization factor (NF) in each sample for quantification of relative expression analyses of the target genes.

**Table 2.** Ranking of reference genes in anterior and posterior scales across all of the haplochromine species used in this study. SD indicates a ranking calculation based on standard deviation generated by BestKeeper, whereas SV, stability value, and M, mean expression stability value, are calculated by geNorm and NormFinder, respectively.

|  | Anterior scales |  |  |  |  |  | Posterior scales |  |  |  |  |  |
| --- | --- | --- | --- | --- | --- | --- | --- | --- | --- | --- | --- | --- |
|  | BestKeeper |  | geNorm |  | NormFinder |  | BestKeeper |  | geNorm |  | NormFinder |  |
|  | Ranks | SD | Ranks | M | Ranks | SV | Ranks | SD | Ranks | M | Ranks | SV |
| 1 | *actb1* | 0.398 | *actb1* | 1.110 | *actb1* | 0.478 | *actb1* | 0.421 | *actb1* | 1.133 | *actb1* | 0.435 |
| 2 | *rps11* | 0.532 | *rps11* | 1.168 | *rps11* | 0.589 | *rps11* | 0.571 | *rps11* | 1.174 | *rps11* | 0.471 |
| 3 | *rps18* | 0.745 | *tbp* | 1.256 | *tbp* | 0.669 | *tbp* | 0.804 | *tbp* | 1.276 | *tbp* | 0.705 |
| 4 | *tbp* | 0.766 | *hsp90a* | 1.379 | *rps18* | 0.708 | *hsp90a* | 0.829 | *hsp90a* | 1.346 | *hsp90a* | 0.763 |
| 5 | *gapdh* | 0.869 | *rps18* | 1.396 | *hprt1* | 0.871 | *rps18* | 0.869 | *rps18* | 1.536 | *rps18* | 0.812 |
| 6 | *hsp90a* | 0.882 | *hprt1* | 1.590 | *hsp90a* | 0.879 | *gapdh* | 1.003 | *gapdh* | 1.703 | *hprt1* | 0.918 |
| 7 | *hprt1* | 0.929 | *gapdh* | 1.631 | *gapdh* | 1.227 | *hprt1* | 1.089 | *hprt1* | 1.729 | *gapdh* | 1.215 |
| 8 | *elf1a* | 1.507 | *elf1a* | 1.827 | *elf1a* | 1.420 | *elf1a* | 1.362 | *elf1a* | 1.754 | *elf1a* | 1.310 |

### Gene expression differences between anterior and posterior scales

The relative expression levels of 16 candidate target genes, *bmp4, col1a2, ctsk, dlx5, eda, edar, fgf20, fgfr1, mmp2, mmp9, opg, rankl, runx2a, sema4d, shh* and *sp7*, were compared between the anterior and posterior scales in each of the haplochromine species (Fig. 3). Some of these genes such as *bmp4, col1a2, rankl* and *sp7*, showed almost no expression difference between the anterior and posterior scales. Moreover, none of the target genes showed consistent expression difference across all the species. These indicate potential involvement of various genes in morphological divergence between the anterior and posterior scales. However, two genes, *ctsk* and *shh* exhibited expression difference between the anterior and posterior scales in most of the species (Fig. 3). The directions of expression differences between the anterior and posterior scales for *ctsk* and *shh* were variable depending on the species. Interestingly all the three species form LT showed higher expression in the anterior scale for *shh*, whereas the all the species from LM and LV showed tendency for opposite pattern with increased posterior scale expression. These findings suggest potential role of *ctsk* and *shh* in morphological divergence of the scales along anterior-posterior axis.

**Figure 3.**
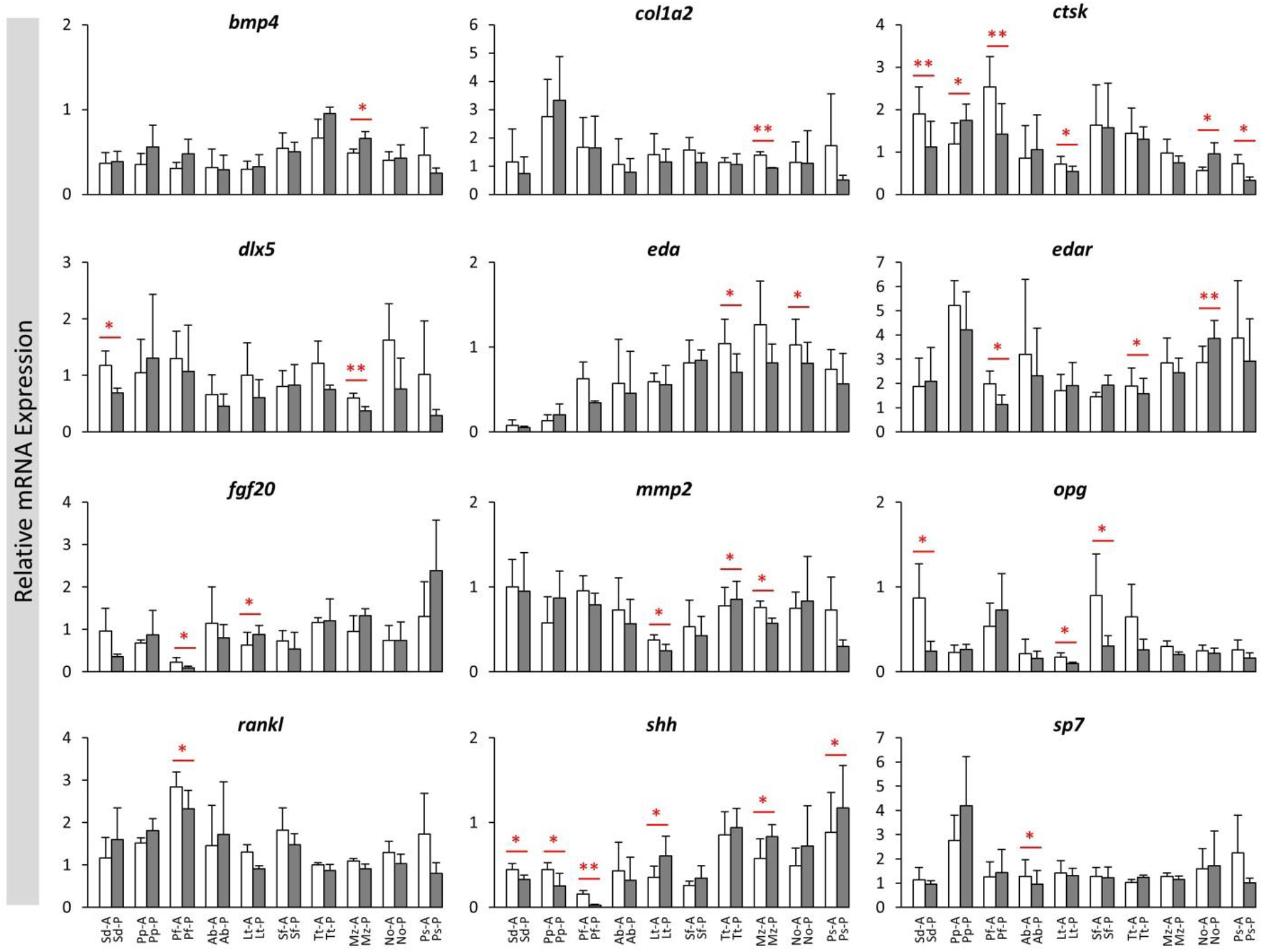
The anterior versus posterior scales expression differences of the candidate target genes in haplochromine cichlids from three East African lakes. Comparisons of relative expression levels between anterior versus posterior scales for 16 candidate target genes in different lakes in East Africa at young adult stage. Significant differences between the are indicated by red asterisks (*P < 0.05; **P < 0.01). See Figure 1A for corresponding species abbreviations.

### Gene expression differences between lakes in anterior and posterior scales

Next, we compared the expression levels of the target genes between the lakes in the anterior or posterior scales by considering all the species from each lake as one group (Fig. 4). In the anterior scales, nine out of 16 target genes showed expression differences between the lakes including *bmp4, ctsk, eda, edar, mmp2, opg, rankl, shh* and *sp7*. Most of these differences were between LT and the other lakes, and 4 genes, *ctsk, mmp2, opg* and *rankl* showed higher expression in LT species, while 3 genes, *bmp4, eda* and *shh* showed lower expression LT species. Furthermore, 2 genes, *ctsk* and *eda* displayed the strongest expression differences between the lakes in opposite patterns suggesting their role in morphological divergence of the anterior scales across the lakes (Fig. 4). In the posterior scales, 11 out of 16 target genes showed expression differences between the lakes including *bmp4, col1a2, ctsk, eda, edar, fgf20, fgfr1, mmp2, opg, rankl, shh* and *sp7*. Again, most of these differences in the posterior scales were between LT and the other lakes, and 5 genes, *col1a2, ctsk, mmp2, opg* and *rankl* showed higher expression in LT species, while 3 genes, *eda, fgf20* and *shh* showed lower expression LT species. In addition, four genes, *fgf20, rankl, eda* and *shh* displayed the strongest expression differences between the lakes in opposite patterns (*eda* and *rankl* higher in LT, and *fgf20* and *shh* lower in LT) suggesting their role in morphological divergence of the posterior scales across the lakes (Fig. 4). In general, more genes with stronger expression differences between the lakes were observed the posterior scales. Several genes such as *ctsk, eda, edar, mmp2, opg, rankl* and *shh* appeared to have similar patterns of expression differences between the lakes in both anterior and posterior scales. Importantly, we found only one gene, *eda*, to have strong differences between the lakes in both anterior and posterior scales indicating its potentially crucial role in morphological divergence of the scales across the lakes.

**Figure 4.**
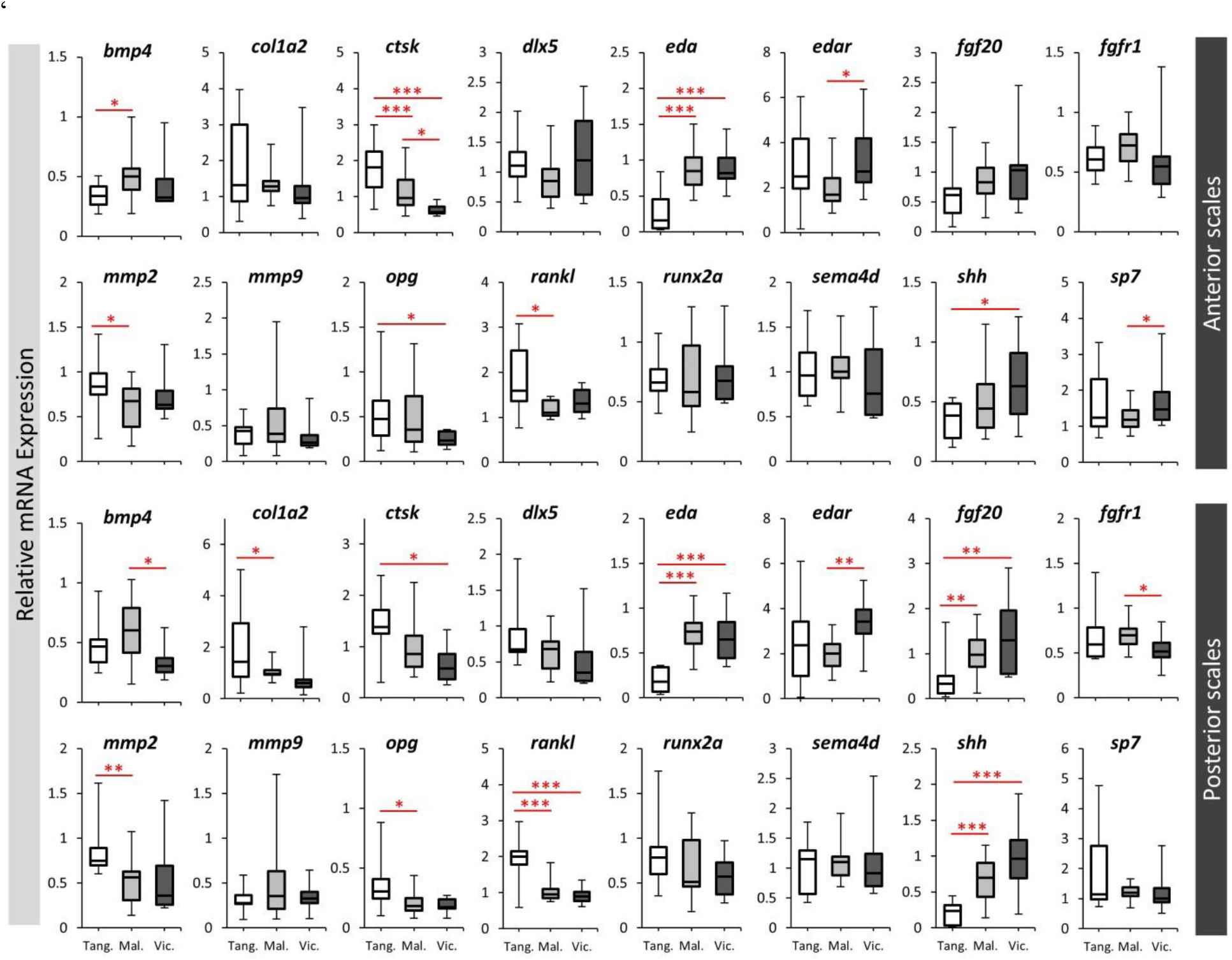
The lake-based expression differences of the candidate target genes in haplochromine cichlids in this study. Comparisons of relative expression levels between the lakes, when all species of each lake were combined, within anterior or posterior scales for 16 candidate target genes. Significant differences between the are indicated by red asterisks (*P < 0.05; **P < 0.01; ***P < 0.001). See Figure 1A for corresponding species abbreviations.

### Correlation analyses between gene expression and morphological divergence in scales

We analysed the correlation between expression of the genes and canonical variate 1 in the anterior or posterior scales across the species. Only one gene, *eda*, showed significant correlation in the anterior scales (Fig. 5), whereas, in the posterior scales 4 genes including *dlx5, eda, rankl* and *shh* displayed significant correlations between expression and morphological differences (Fig. 6). Among these genes *eda* exhibited the strongest correlation in the posterior scales. However, the correlation patterns differed between the genes in the posterior scales, i.e. eda and shh showed positive while *dlx5* and *rankl* had negative correlations with the morphological changes based canonical variate 1. Therefore, again only one gene, *eda*, showed significant correlation between its expression and the morphological differences in both scales indicating its potential role in divergent scale morphogenesis in the cichlid species. The opposite correlation patterns in the posterior scales might also indicate inhibitory regulatory connections between the genes.

**Figure 5.**
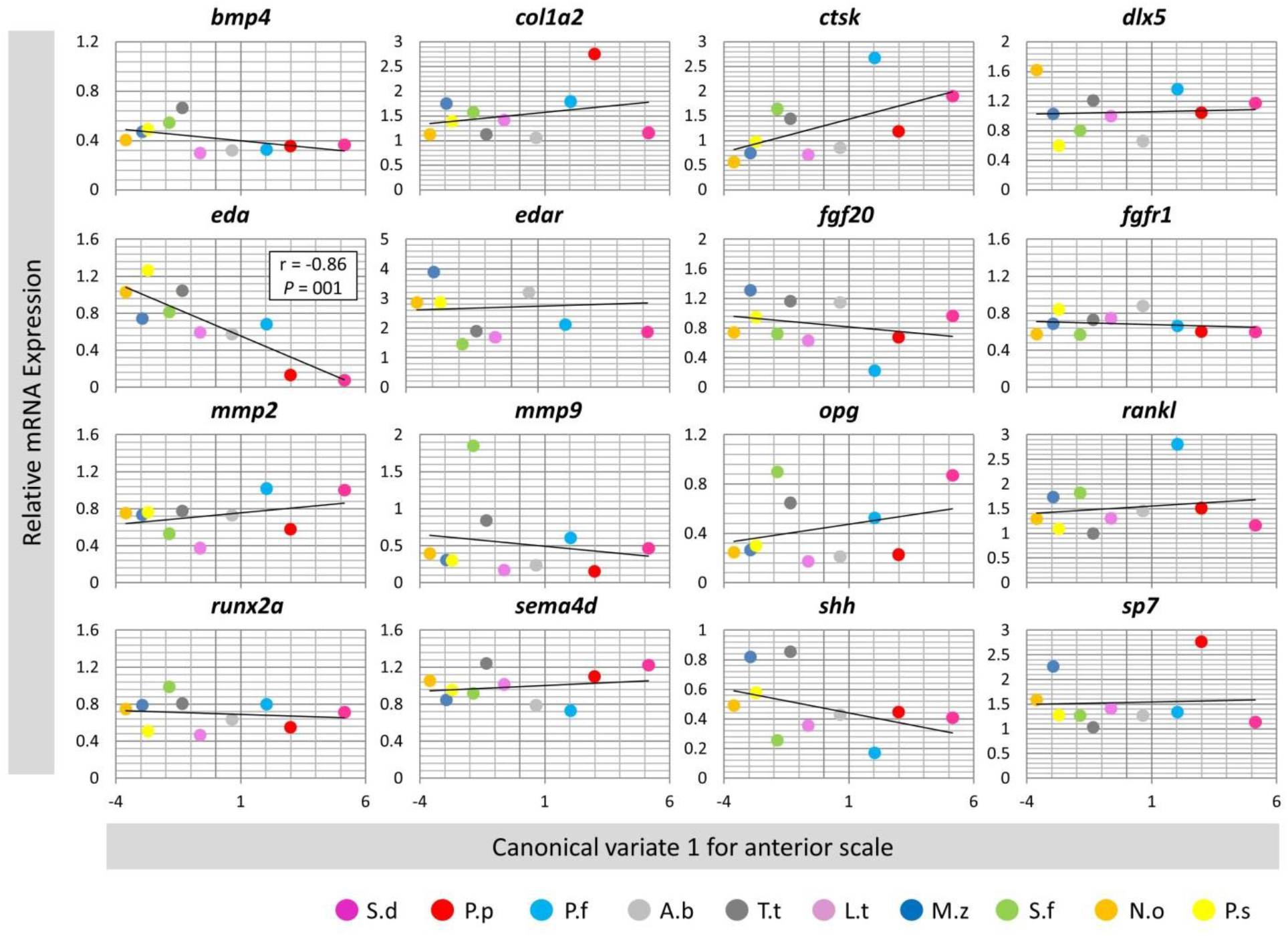
Correlation analyses of candidate target gene expressions and the anterior scale morphololgical divergence across the haplochromine species. A Pearson correlation coefficient (r) was used to assess the similarity between differences in expression level of the target genes and the major canical variate in the anterior scales across all species. See Figure 1A for corresponding species abbreviations.

**Figure 6.**
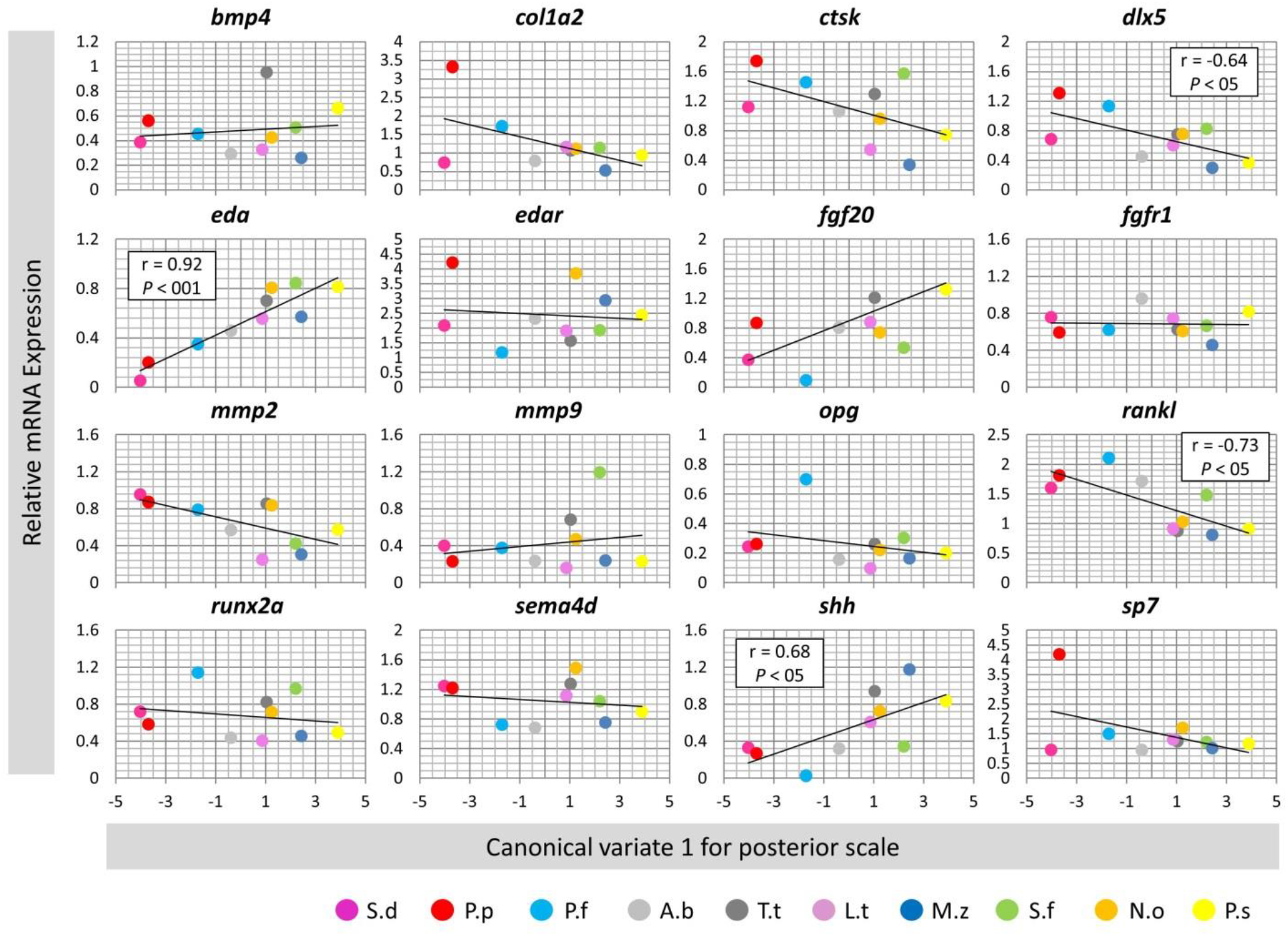
Correlation analyses of candidate target gene expressions and the posterior scale morphololgical divergence across the haplochromine species. A Pearson correlation coefficient (r) was used to assess the similarity between differences in expression level of the target genes and the major canical variate in the posterior scales across all species. See Figure 1A for corresponding species abbreviations.

### Genetic differences in non-coding sequences of *eda* gene

Finally, we were interested to investigate genetic differences in available regulatory sequences of *eda* gene including 5’UTR, 3’UTR and short but conserved inter-genic region between *eda* and *tnfsf13b* (immediate downstream gene) across the species. Interestingly, we found two genetic differences (mutations/deletions) in 3’UTR and one in *eda*-*tnfsf13b* intergenic region to differ between LT species versus LM and LV species (Table 3). Next, we parsed the short sequence regions containing the mutations/deletions against transcription factor binding site (TFBS) databases. We found that the two changes in 3’UTR seem to lead to gaining TFBS for transcription factors Mef2 and Tcf1 in the LM and LV species, whereas the changes in the inter-genic region led to gaining TFBS for Lef1 transcription factor in the LT species (Table 3). Importantly, the riverine species A.b appeared to have intermediate genetic changes meaning that for the two changes in 3’UTR it showed a deletion similar to the LT species but a mutation similar to the LV and LM species. Also, for the inter-genic change, A.b showed an intermediate mutation between the LT and the other species from LM and LV, however, this mutation showed no gain of TFBS (similar to the LM and LV species). Taken together, these genetic changes showed similarity with differences in gene expression and scale morphology, where the LT species clustered different from LM and LV species and the riverine species (A.b.) showed intermediate differences. This suggests that the identified genetic changes might be the underlying factors for divergent *eda* expression as well as differences in the scale morphology.

**Table 3.**
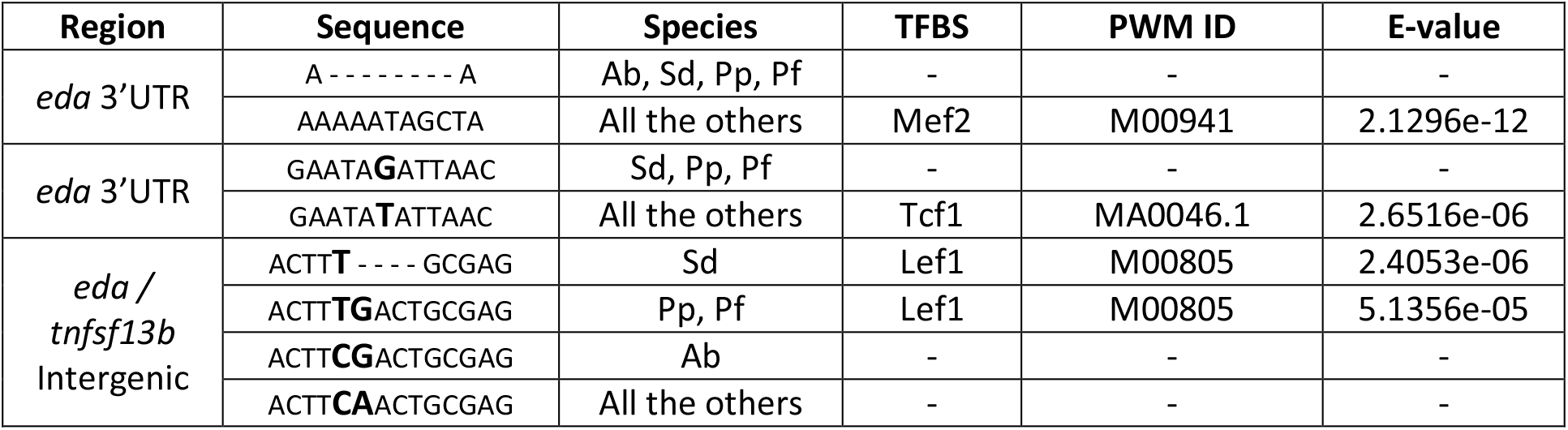
Identified genetic differences in regulatory sequences of *eda* gene and predicted binding sites for potential upstream regulators. PWD ID indicates positional weight matrix ID of a predicted binding site and E-values refer to matching similarity between the predicted motif sequences and the PWD IDs.

| Region | Sequence | Species | TFBS | PWM ID | E-value |
| --- | --- | --- | --- | --- | --- |
| *eda* 3'UTR | A - - - - - A | Ab, Sd, Pp, Pf | - | - | - |
|  | AAAAATAGCTA | All the others | Mef2 | M00941 | 2.1296e-12 |
| *eda* 3'UTR | GAATA**G**ATTAAC | Sd, Pp, Pf | - | - | - |
|  | GAATA**T**ATTAAC | All the others | Tcf1 | MA0046.1 | 2.6516e-06 |
| *eda* /<br>*tnfsf13b*<br>Intergenic | ACTT**T** - - - - GCGAG | Sd | Lef1 | M00805 | 2.4053e-06 |
|  | ACTT**TG**ACTGCGAG | Pp, Pf | Lef1 | M00805 | 5.1356e-05 |
|  | ACTT**CG**ACTGCGAG | Ab | - | - | - |
|  | ACTT**CA**ACTGCGAG | All the others | - | - | - |

## Discussion

As river-adapted haplochromine cichlids repeatedly seeded adaptive radiations in several East African lakes, cichlid fishes recurrently adapted to corresponding trophic niches. Thereby, the adaptive value of traits is often mirrored by morphological shape parallelism and concomitant similar lifestyles which result from parallel evolution [4, 5]. Hence, fishes from the cichlid species flocks in various African lakes comprise an exciting model to conduct comparative morphological and molecular studies. While most previous studies focused on bony elements that can easily be linked particular trophic niches and divergent natural selection as driver of diversification in cichlids (e.g., [52]), other skeletal structures such as scales might show less obvious adaptive trajectories.

Above all, between the three East African Great Lakes, the haplochromine cichlids are especially interesting, as they share common ancestry and comprise the Tropheini at LT and the entire the LV and LM haplochromine radiations [53, 54]. As the lakes have all very different geological histories, with Lake Tanganyika being the oldest [55], LM the intermediate [56] and LV the youngest of the three [57], they also depict three extensive radiations at different time points. Thus, depending on the evolutionary age of the different lakes, species (and their morphologies) had more or less time to diverge, despite sharing parallels. The more time passes, much more elaborated predator-prey and host-parasite relationships can evolve. This is manifested by unique ecological and behavioural features, particularly in the oldest of the three lakes, LT, which contains coocoo-catfish species showing brood parasitism [58], dwarfed gastropod shell breeders [59], putative cleaning behaviour [60], or highly elaborated scale eaters [61] (but also see *Genyochromis mento* from Lake Malawi). Particularly the latter case, scale eating, could have influenced the co-evolution of scale morphology in host species. Lake Tanganyika’s scale eaters (i.e., *Perissodus*; Perissodini) show different degrees of specialization, whereas only the shallow water species, *Perrisodus microlepis* and *P. straeleni*, feed almost exclusively on fish scales while other species are not that specialized [4, 61]. The most common prey species of *P. microlepis* are members of the Tropheini and Eretmodini [62]. *P. straeleni* seems to be less specialised to certain prey items, but Tropheini scales still make up a major part of the gut contents [63]. Based on our dataset (Tropheini only) it remains speculative to assume that scale-related gene expression and the concomitant morphology might reflect an adaptation to reduce the risk of scale predation. Future studies, including more early branching (non-modern) haplochromine cichlids (e.g., *Pseudocrenilabrus, Thoracochromis, Astatoreochromis*) or other Tropheini with different lifestyles (e.g., *Ctenochromis*) will be necessary to establish stronger links between scale morphology and this unique predation pressure.

Nonetheless, understanding which genetic mechanisms underlie the scale morphology might be the key to understand how such similar and/or divergent eco-morphologies evolved. Perhaps the most striking finding of our study was the highly significant differential expression of *eda* between LT species versus the species from the younger lakes (LM and LV species) in both anterior and posterior scales (Fig. 4). Interestingly, the *eda* expression in the riverine species (Ab), which is believed to be an ancestral species to haplochromine cichlids of the three great African lakes, was at intermediate level between LT and the species from LM and LV in both anterior and posterior scales. Moreover, the expression patterns of *eda* in both scales were highly correlated with morphological divergence across the species in this study. Ectodysplasin A (*eda*) encodes a member of the tumor necrosis factor family and mediates a signal conserved across vertebrates which is essential for morphogenesis of ectodermal appendages, such as scale, hair and feathers [22]. The eda signal is mediated through its receptor (encoded by *edar*) and initiated upon binding of eda to edar on the surface of a target cell [22]. In human, mutations in components of eda signal can cause hypohidrotic ectodermal dysplasia (HED) which is characterized by reduction and abnormal teeth morphology, absence or reduction of sweating glands and hair [64]. Similarly, impaired eda signal in zebrafish and medaka can lead to reduction in the number of scales and teeth [18, 19]. In sculpin (*Cottus*) fishes, genetic changes in the receptor gene (*edar*) has been found to be associated with morphological variations in body prickles (calcified spicules embedded in the skin), which are homologous structures to fish scales [65]. A later study in a highly derived order of teleosts, Tetraodontiformes, which includes ocean sunfishes, triggerfishes and pufferfishes, also showed the importance of eda signaling pathway in developmental formation and morphological variations of dermal spines (an extreme scale derivative) [66]. In stickleback, a mutation within an inter-genic region between *eda* and *tnfsf13b* genes leads to changes in transcriptional responsiveness of *eda* to its upstream Wnt signaling pathway and consequently impairment of armor plate formation [67].

In this study, we also found genetic changes in 3’UTR and the inter-genic region between *eda* and *tnfsf13b* genes that could explain the differences in *eda* expression across the Haplochromine cichlids (Table 3). The genetic changes resulted gain or loss of motifs which were predicted to be binding sites for transcription factors encoded by *mef2, tcf1* and *lef1* genes. These changes always discriminated the LT species from the species from LM and LV, whereas the riverine species had changes which could be considered an intermediate to both groups. Interestingly, all of the three predicted transcription factors (*mef2, tcf1* and *lef1*) are linked to Wnt signaling pathway. It is already known that mef2 can enhance canonical Wnt signal [68] and it is involved in osteogenesis as well [69–71]. The binding site motif for mef2 appeared to be deleted in 3’UTR of the LT and riverine species. On the other hand, a binding site motif for tcf1 was gained in 3’UTR of the LM, LV and riverine species. In mice, *tcf1* is demonstrated to be involve in paraxial mesoderm and limb formation and appeared to be act downstream of Wnt signal similar to lef1 transcription factor [72]. Moreover, canonical Wnt signaling has been shown to regulate osteogenesis through tcf1 responsive element on regulatory sequence of runx2 in mammals [73]. The third motif predicted to be a binding site for lef1 and only found within *eda* - *tnfsf13b* inter-genic region of LT species. Lef1 is again a well-known mediator of canonical Wnt signaling pathway which inhibits final stage of osteoblast differentiation [74] but it is essential for osteoblast proliferation and normal skeletal development [75, 76]. During development lef1 function is shown to be essential for scale outgrowth in zebrafish [77], and *eda* expression is known to be regulated by Wnt signal through *lef1* transcriptional activity in mammals [78, 79]. In stickleback, mutation in an inter-genic region between *eda* and *tnfsf13b* genes is suggested to affect a binding site for c-jun transcription factor which its interaction with lef1 is required for *eda* transcriptional response to Wnt signal during armor plate formation [67]. Taken together, these observations, suggest mutations in enhancer sequences required for binding of Wnt signal components as potential underlying reason for the divergent expression of *eda* in both anterior and posterior scales of the cichlid species in this study.

In the posterior scale, in addition to *eda*, three more genes, *dlx5, rankl* and *shh*, displayed expression correlation with morphological divergence across the cichlid species (Fig. 6). The first gene, distal less homeobox 5 or *dlx5*, encodes transcription factor stimulating osteoblast differentiation and bone development, and it is also implicated in scale development and regeneration in fish [80–82]. Apart from its role in skeletogenesis, *dlx5* has been found to be involved in divergent development and morphogenesis of other tissues in cichlids such as teeth and nuchal hump [39, 83, 84]. In goldfish, *dlx5* expression appeared to be important at early stages of scale regeneration [80], and in both zebrafish and goldfish, *dlx5* transcription in scale can be affected by environmental clues such as mechanical stimulus [81, 82]. The second gene, *rankl*, encodes a ligand for osteoprotegerin (opg) and play crucial role in osteoclast differentiation and bone remodelling. Changes in *rankl* transcription appeared to be important during scale regeneration in goldfish [80, 85], as well as intercellular communications regulating scale bone remodeling in zebrafish and goldfish [81, 86]. Both *dlx5* and *rankl* have shown expression correlation patterns opposite to *eda* and *shh* in the posterior scales. Although, direct regulatory connections between these factors have not been investigated in scale but these findings suggest their potential interactions at transcriptional level. Moreover, higher expression of *rankl* in the scales of LT species compared to LM and LV species might indicate higher level of bone remodelling in their scales.

The third gene, sonic hedgehog or *shh*, encodes a ligand of hedgehog signaling pathway which is shown to control scale morphogenesis in relationship with the formation of the epidermal fold in the posterior region of scale in fish [16]. In zebrafish, epidermal expression of *shh* has been shown to regulate scale regeneration through controlling osteoblast population and affecting directional bone growth [87]. We found similar expression pattern between *eda* and *shh* which is more pronounced in the posterior scales. This is consistent with previous findings in other vertebrates, for instance, *eda* has been demonstrated to act upstream of *shh* and induce *shh* expression during ectodermal organogenesis in mammals (e.g. during hair placode formation) [88–92]. Furthermore, it has been shown that the *eda*-dependent regulation of *shh* might be a part of larger molecular cascade in which an upstream signal such as Wnt pathway activates *eda* signal and in turn *eda* induces *shh* transcription [90, 92, 93]. These observations suggest potential role of *Wnt*-*eda*-*shh* axis in divergent scale morphogenesis across Haplochromine cichlids, which seems to be more pronounced in the posterior scales.

## Conclusions

This is the first attempt to study cross-species association between gene expression and morphological divergence in scales of cichlids from different lakes. Our results provide evidence for potential role of a key signal mediated by *eda* gene to be involved in divergent morphogenesis of scale in closely related cichlid species. We show that *eda* expression has lower level in the scales of species from the older lake (Lake Tanganyika) and correlates with the observed shape variations across species. Our findings shed light on molecular basis of morphological divergence of a less studied skeletal element; however, further investigations are required to understand whether these differences have adaptive relevance in ecological and evolutionary-developmental contexts.

## List of abbreviations

LT: Lake Tanganyika
LM: Lake Malawi
LV: Lake Victoria
*bmp4*: bone morphogenetic protein 4
*col1a2*: collagen type I alpha 2 chain
*ctsk*: cathepsin K
*dlx5*: distal-less homeobox 5
*eda*: ectodysplasin A
*edar*: ectodysplasin A receptor
*fgf20*: fibroblast growth factor 20
*fgfr1*: fibroblast growth factor receptor 1
*mmp2*: matrix metallopeptidase 2
*mmp9*: matrix metallopeptidase 9
*opg*: osteoprotegerin
*rankl*: receptor activator of nuclear factor kappa B ligand
*runx2a*: runt-related transcription factor 2 alpha
*sema4d*: semaphorin 4D
*shh*: sonic hedgehog signaling molecule
*sp7*: osterix transcription factor.

## Declarations

### Authors’ contributions

EPA, SB, MW and CS designed the study. SB, EPA, MW, and AD conducted the laboratory experiment, measurements and figure preparations. MW and EPA analysed the data, and EPA, MW, AD and CS wrote the manuscript. WG and AD performed fish breeding and sampling. WG photographed the adult fishes used in Figure 1A. All authors reviewed the manuscript and approved its content.

## Acknowledgements

The authors acknowledge Institute of Biology at University of Graz for providing fish breeding and laboratory facilities, and the Austrian Science Fund for the financial support. We also thank Stephan Koblmüller for sharing his precious knowledge on cichlid fishes of the African Lakes.

## Availability of data and materials

All data generated or analysed during this study are included in this published article.

## Consent for publication

Not applicable.

## Ethics approval and consent to participate

Fish keeping and euthanasia were conducted under permit BMWFW-66.007/0004-WF/V/3b/2016 issued by the Federal Ministry of Science, Research and Economy of Austria (BMWFW) in accordance with the ethical guidelines and regulations of the BMWFW. Fish keeping and sampling was carried out in our certified aquarium facility according to the Austrian animal welfare law.

---

**Additional file 1.xls.** Information about qPCR primers used in this study.

